# Origin, Conservation, and Loss of Alternative Splicing Events that Diversify the Proteome in Saccharomycotina Budding Yeasts

**DOI:** 10.1101/2020.04.13.039594

**Authors:** Jennifer E. Hurtig, Minseon Kim, Luisa J. Orlando-Coronel, Jellisa Ewan, Michelle Foreman, Lee-Ann Notice, Michelle A. Steiger, Ambro van Hoof

## Abstract

Many eukaryotes use alternative splicing to express multiple proteins from the same gene. However, while the majority of mammalian genes are alternatively spliced, other eukaryotes use this process less frequently. The budding yeast *Saccharomyces cerevisiae* has been successfully used to study the mechanism of splicing and the splicing machinery, but alternative splicing in yeast is relatively rare and has not been extensively studied. We have recently shown that the alternative splicing of *SKI7/HBS1* is widely conserved, but that yeast and a few other eukaryotes have replaced this one alternatively spliced gene with a pair of duplicated unspliced genes as part of a whole genome doubling (WGD). Here we show that other examples of alternative splicing that were previously found to have functional consequences are widely conserved within the Saccharomycotina. We also show that the most common mechanism by which alternative splicing has disappeared is by the replacement of an alternatively spliced gene with duplicate genes. Saccharomycetaceae that diverged before WGD use alternative splicing more frequently than *S. cerevisiae*. This suggests that the WGD is a major reason for the low frequency of alternative splicing in yeast. We anticipate that whole genome doublings in other lineages may have had the same effect.

## Introduction

Alternative splicing is thought to be an important mechanism to generate multiple functionally distinct proteins from a single gene. Although the budding yeast *Saccharomyces cerevisiae* has proven to be a powerful tool to understand the mechanism of splicing, only three genes are known to encode two functionally distinct proteins through alternative splicing. Both *PTC7* and *FES1* can be spliced or the intron can be retained, and in both cases this results in altered localization of the protein (Gowda et al., 2016; Juneau et al., 2009). Fes1 is a nucleotide exchange factor for Hsp70 that is targeted to the nucleus when translated from the spliced mRNA and remains cytoplasmic when translated from the unspliced mRNA (Gowda et al., 2016). Similarly, Ptc7 is a protein phosphatase that is localized inside the mitochondria when translated from the spliced mRNA. Translation of the unspliced *PTC7* mRNA result in the inclusion of a transmembrane helix and insertion of Ptc7 into the nuclear membrane (Juneau et al., 2009). In contrast to the well-characterized effects on Ptc7 and Fes1 localization, the functional consequences of *SRC1* alternative splicing are only partially understood. *SRC1* generates two proteins through the use of alternative 5’ splice sites (Grund et al., 2008; Rodriguez-Navarro et al., 2002). The alternative 5’ splice sites of *SRC1* are separated by 4 nucleotides, resulting in the downstream exon being used in two different reading frames. Both the 95 KDa “full-length” and 73 KDa “truncated” proteins are inserted into the nuclear membrane. The lack of the full-length protein causes growth defect in mutants with defect in the THO complex, suggesting that the full-length protein has some function that is not shared with the truncated protein (Grund et al., 2008). The full-length protein is also similar to a WGD paralog, Heh2, and the functional differences between Src1 and Heh2 are also not well understood.

We have recently shown that in the related yeast *Lachancea kluyveri* a single gene encodes Ski7 and Hbs1 through the use of alternative splicing, but that this alternatively spliced gene is replaced by separate *SKI7* and *HBS1* genes in *S. cerevisiae* (Marshall et al., 2013). Ski7 tethers the Ski complex to the cytoplasmic RNA exosome complex and thereby mediates mRNA degradation (Araki et al., 2001; Kowalinski et al., 2016; van Hoof et al., 2000). Hbs1 instead delivers Dom34 to stalled ribosomes, which allows Dom34 to recycle the ribosomal subunits for subsequent rounds of translation (Becker et al., 2011; Pisareva et al., 2011; Shoemaker et al., 2010). We further showed that this alternative splicing of *SKI7/HBS1* is conserved in animals, fungi, and plants (Marshall et al., 2018; Marshall et al., 2013).

During our studies focused on *SKI7/HBS1*, we noted anecdotally that two other gene pairs fit the same pattern. First, the *S. cerevisiae YSH1* and *SYC1* genes arose by duplication of an alternatively spliced *YSH1/SYC1* gene that uses alternative 3’ splice sites (Marshall et al., 2018). In *S. cerevisiae*, Ysh1 is the endonuclease that is responsible for the 3’end formation of mRNAs in the cleavage and polyadenylation reaction, while Syc1 is required for cleavage-independent 3’ end formation of some noncoding RNAs (Lidschreiber et al., 2018). Syc1 shares some protein-protein interactions with Ysh1, but lacks the endonuclease domain (Nedea et al., 2003). In *L. kluyveri*, RNA-seq indicates that the Ysh1 and Syc1proteins are encoded by a single gene through the use of alternative 3’ splice sites (Marshall et al., 2018). The other previously noted example of an alternatively spliced gene being replaced by duplicate genes is *PTC7*. As noted above, the *S. cerevisiae PTC7* gene encodes two protein phosphatases with distinct localization. (Juneau et al., 2009). In contrast, *Tetrapisispora blattae* contains two *PTC7* genes, and we have speculated that one contains an intron. This predicted intron contains stop codons that prevent translation of any intron-retained mRNA (Marshall et al., 2013). However, there is no experimental evidence supporting this suggestion and introns in *T. blattae* have not been carefully annotated. Unlike *SKI7/HBS1* alternative splicing, which is conserved in animals, fungi and plants, neither the *YSH1/SYC1* or *PTC7* alternative splicing events appeared to be conserved from an ancient ancestor, but when these alternative splicing events arose has not been extensively studied.

Strikingly, the duplication of *SKI7* and *HBS1, YSH1* and *SYC1* and *PTC7* each date to a whole genome doubling (WGD) that occurred an estimated 95 million years ago (MYA) (Kellis et al., 2004; Marcet-Houben and Gabaldon, 2015; Scannell et al., 2007). This WGD also gave rise to the *SRC1* and *HEH2* paralogs mentioned above. These observations suggest that the WGD event in the *Saccharomyces* lineage may be a contributor to the relatively infrequent use of alternative splicing to diversify its proteome, but the relationship between WGD and loss of alternative splicing is incompletely understood. Here, we more systematically analyze when previously identified examples of alternative splicing in Saccharomycotina arose, how extensively they are conserved, whether they were replaced by duplicated genes, and whether, when, and how alternative splicing mechanisms have changed.

## Results and Discussion

### Study design

The subphylum Saccharomycotina is divided into approximately a dozen clades, which generally correspond to families, and in the current study we included all the clades/families with available genome and RNA-seq data ((Shen et al., 2016); Figure 1). We sampled the Saccharomycetaceae family more densely to gain a better understanding of the relationship between WGD and loss of alternative splicing. The Saccharomycetaceae are divided into three clades: the KLE, ZT, and WGD clades (Marcet-Houben and Gabaldon, 2015; Shen et al., 2016). The KLE clade is composed of the *Kluyveromyces, Lachancea* and *Eremothecium* genera, while the ZT clade is composed of the *Zygosaccharomyces* and *Torulaspora* genera. The WGD clade arose through a single hybridization between one parent from the KLE clade and another parent from the ZT clade (Marcet-Houben and Gabaldon, 2015). After this single hybridization, the WGD clade diverged into *Saccharomyces, Kazachstania, Naumovozyma, Tetrapisispora* and *Vanderwaltozyma* genera. The original hybridization occurred before the extant species of the KLE and ZT clades diverged from each other and thus the parents can not be meaningfully be assigned to one of the extant species. We therefore included two diverse representatives from both the KLE and ZT clades, and four from the WGD clade. Within each clade we maximized the diversity analyzed. For example, the first branch within the WGD clade separates the *T. blattae* lineage from the *S. cerevisiae* lineage and we therefore included both of these species.

**Figure 1:**
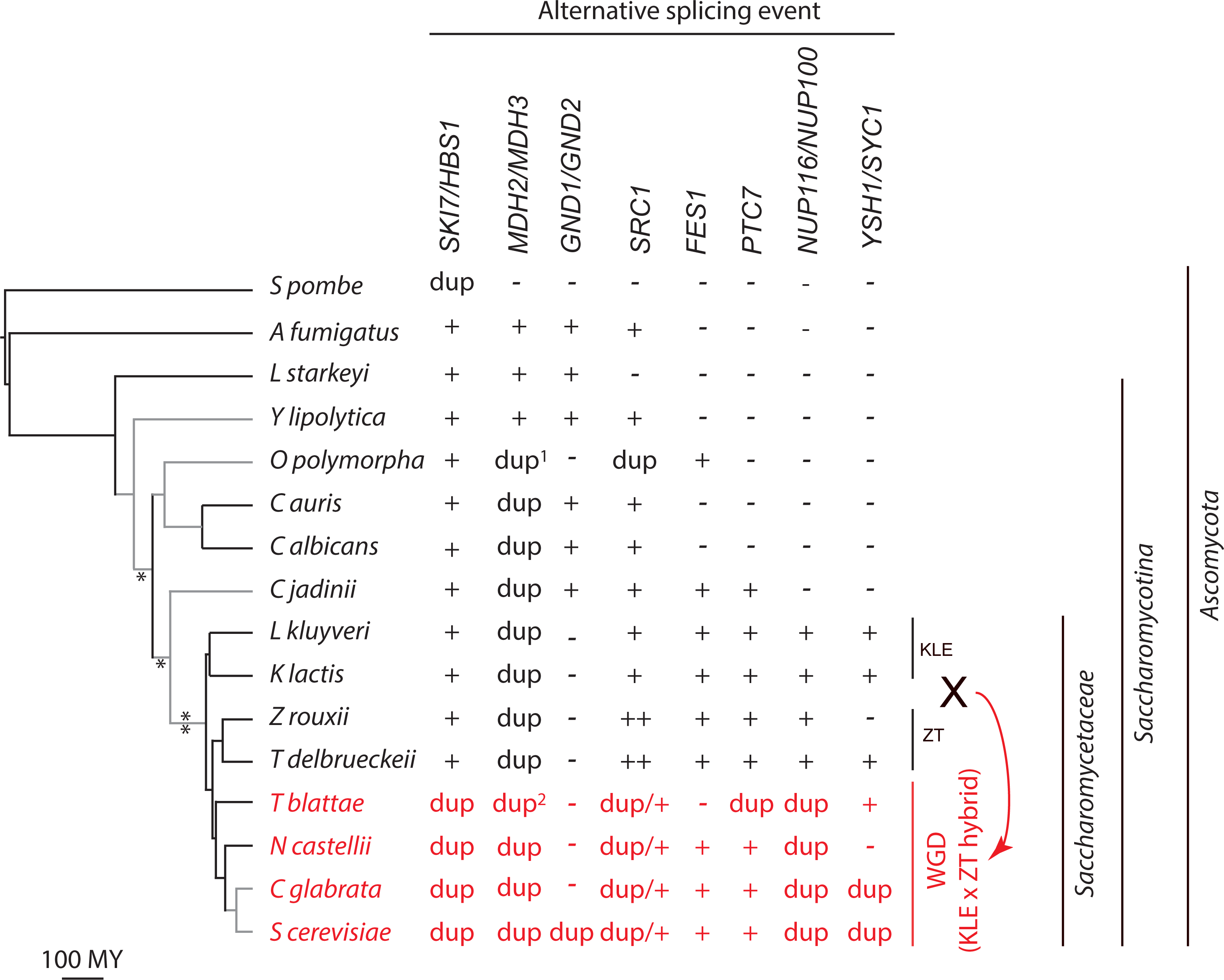
Origin, conservation and loss of alternative splicing events in the Saccharomycotina. Alternative splicing detected in RNA-seq data analysis is indicated by +. Absence of alternative splicing is indicated by -. dup indicates that an alternatively spliced gene was replaced with duplicated genes. The ++ for the ZT clade indicates that the *SRC1* gene uses both intron retention and alternative 5’ splice sites. ^1^: After duplication the *OGAPODRAFT_15721* gene evolved into a pseudogene and thus the species appears to have lost peroxisomal *MDH*. ^2^: The *T. blattae MDH2* gene for cytoplasmic MDH was retained as duplicates after WGD (*TBLA_0B08630* and *TBLA_0A07800*), in addition to the earlier duplication that resulted in cytoplasmic *MDH2* and peroxisomal *MDH3*. The WGD resulting from hybridization between the KLE clade and ZT clade is highlighted in red. Species branch order is from Shen et al. (2016) with branch lengths adjusted to reflect approximate date of divergence from timetree.org. For the branches in grey timetree.org does not have a divergence time. A * next to a branch indicates that alternative splicing arose in a clade within the Saccharomycotina. Full species names are included in Extended Table 1.

To find alternatively spliced target genes we performed extensive literature searches for “alternative splicing” and each species name. We included only alternative splicing events with some evidence that the alternative splicing event resulted in functionally distinct proteins (to eliminate splicing errors and heterogeneity in splicing without functional consequences). This literature search was supplemented by inspecting RNA-seq data from two species, *L. kluyveri* and *Candida albicans*, which identified one novel intron retention event (in the *L. kluyveri NUP116/NUP100* gene). This resulted in eight alternative splicing events that had clear functional consequences (summarized in Table 1).

**Table 1:**
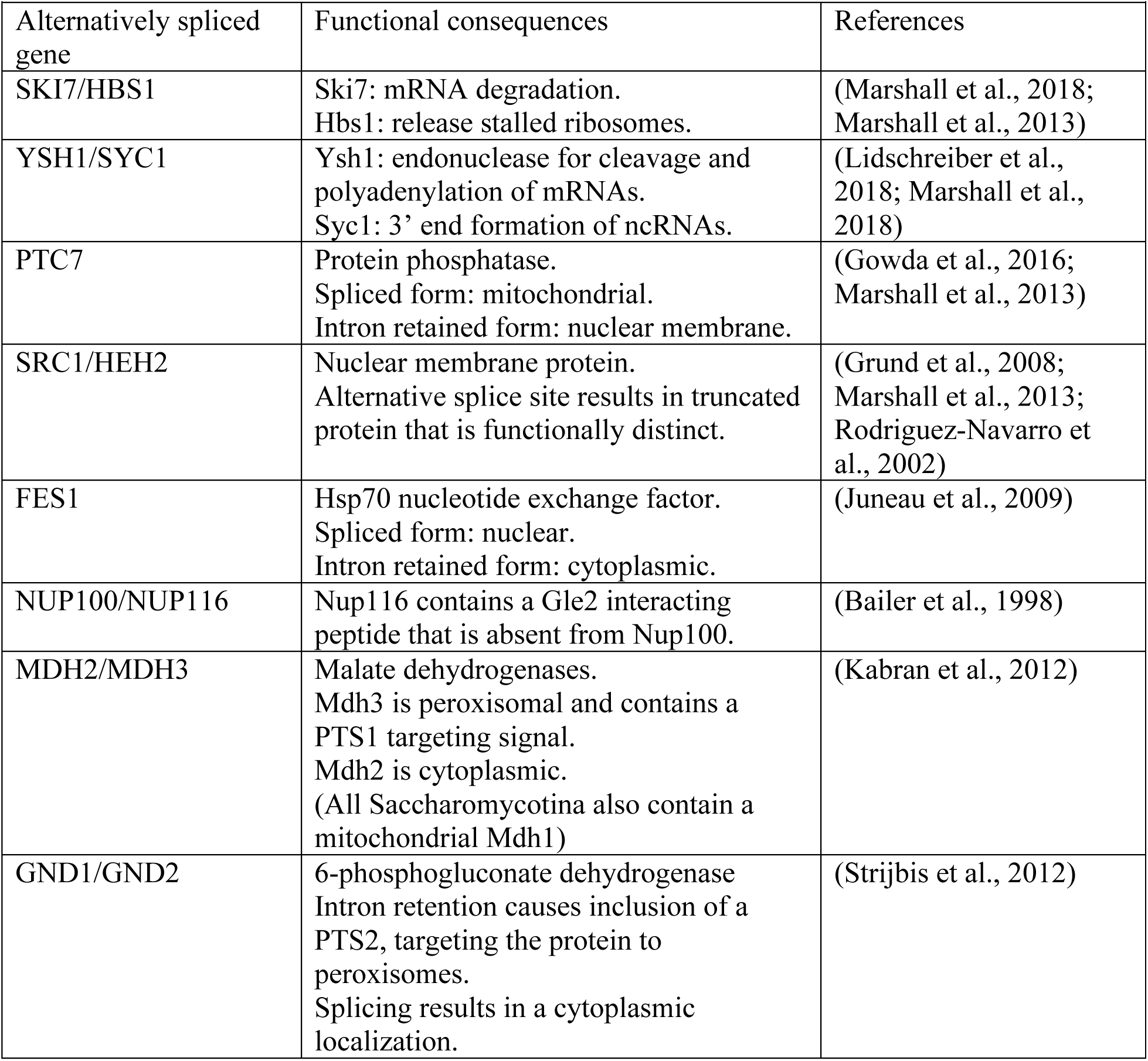
Alternatively spliced genes in Saccharomycotina that result in functional diversification of the proteome

|  |  |  |
| --- | --- | --- |
| Alternatively spliced gene | Functional consequences | References |
| SKI7/HBS1 | Ski7: mRNA degradation.<br>Hbs1: release stalled ribosomes. | (Marshall et al., 2018; Marshall et al., 2013) |
| YSH1/SYC1 | Ysh1: endonuclease for cleavage and polyadenylation of mRNAs.<br>Syc1: 3' end formation of ncRNAs. | (Lidschreiber et al., 2018; Marshall et al., 2018) |
| PTC7 | Protein phosphatase.<br>Spliced form: mitochondrial.<br>Intron retained form: nuclear membrane. | (Gowda et al., 2016; Marshall et al., 2013) |
| SRC1/HEH2 | Nuclear membrane protein.<br>Alternative splice site results in truncated protein that is functionally distinct. | (Grund et al., 2008; Marshall et al., 2013; Rodriguez-Navarro et al., 2002) |
| FES1 | Hsp70 nucleotide exchange factor.<br>Spliced form: nuclear.<br>Intron retained form: cytoplasmic. | (Juneau et al., 2009) |
| NUP100/NUP116 | Nup116 contains a Gle2 interacting peptide that is absent from Nup100. | (Bailer et al., 1998) |
| MDH2/MDH3 | Malate dehydrogenases.<br>Mdh3 is peroxisomal and contains a PTS1 targeting signal.<br>Mdh2 is cytoplasmic.<br>(All Saccharomycotina also contain a mitochondrial Mdh1) | (Kabran et al., 2012) |
| GND1/GND2 | 6-phosphogluconate dehydrogenase<br>Intron retention causes inclusion of a PTS2, targeting the protein to peroxisomes.<br>Splicing results in a cytoplasmic localization. | (Strijbis et al., 2012) |

We then mapped RNA-seq reads from each of the target species to the genome sequence and inspected each of the genes of interest for evidence of alternative splicing. Although alternative splicing is notoriously difficult to annotate *de novo*, the previously described events were obvious in RNA-seq alignments. The use of alternative splice sites was reflected in exon-junction reads where the read starts in one exon and ends in the next exon, precisely defining the splice sites used. Thus, RNA-seq analysis reliably confirmed each of the previously reported alternative splicing events from Table 1.

### Origins and conservation of alternative splicing events in Saccharomycotina

As expected from our earlier studies (Marshall et al., 2018; Marshall et al., 2013), we detected the use of alternative *SKI7/HBS1* 3’ splice sites in early diverging Saccharomycotina. In addition, we detected alternative splicing of three other genes in the early diverging species *Lipomyces starkeyi* and/or *Yarrowia lipolytica* (*MDH2/MDH3, GND1/GND2*, and *SRC1/HEH2*; Figure 1). This suggests that alternative splicing was already present in the common ancestor of the Saccharomycotina. To confirm this, we analyzed alternative splicing in RNA-seq data from *Aspergillus fumigatus* and *Schizosaccharomyces pombe*, which are representatives of the two other major clades of Ascomycota. Consistent with the *L. starkeyi*, and *Y. lipolytica* findings, all four genes were also alternatively spliced in *A. fumigatus*. We conclude that the *SKI7/HBS1, MDH2/MDH3, SRC1/HEH2* and *GND1/GND2* alternative splicing events arose over 500 million years ago (MYA) and have been conserved since then.

Additional alternative splicing events apparently arose later (Figure 2). Specifically, we detected intron retention of *FES1* in *Ogataea polymorpha, Cyberlindnera jadinii*, and Saccharomycetaceae (Figure 2 B to D). In each case the retained intron is in the exact same position in the gene. In *S. cerevisiae* the second exon of *FES1* has been shown to add a 16 amino acid Lys and Arg rich sequence that functions as a Nuclear Localization Sequence (Gowda et al., 2016). In these other species the second exon is similarly short (14-17 AA) and Lys/Arg-rich. Alternatively, when the intron is retained translation stops four to seven codons into the intron. Thus, *FES1* intron retention is clearly conserved in these species, and arose before *Ogataea* diverged from *Saccharomyces* 304 (MYA). Similarly, intron retention in *PTC7* was clearly detected in other Saccharomycetaceae and in *Cyberlindnera jadinii*. The time of divergence of *C. jadinii* has not been established, but *PTC7* intron retention most likely arose after *C. albicans* diverged 304 MYA but before the Saccharomycetaceae diverged from each other 114 MYA. The Saccharomycetaceae contain an additional intron retention event in *NUP116/NUP100* (Figure 2 E to G) and alternative 3’ splice sites in the *YSH1/SYC1* gene (Figure 2 H to J) that also likely arose more than 114 MYA, but are not shared with *C. jadinii*.

**Figure 2:**
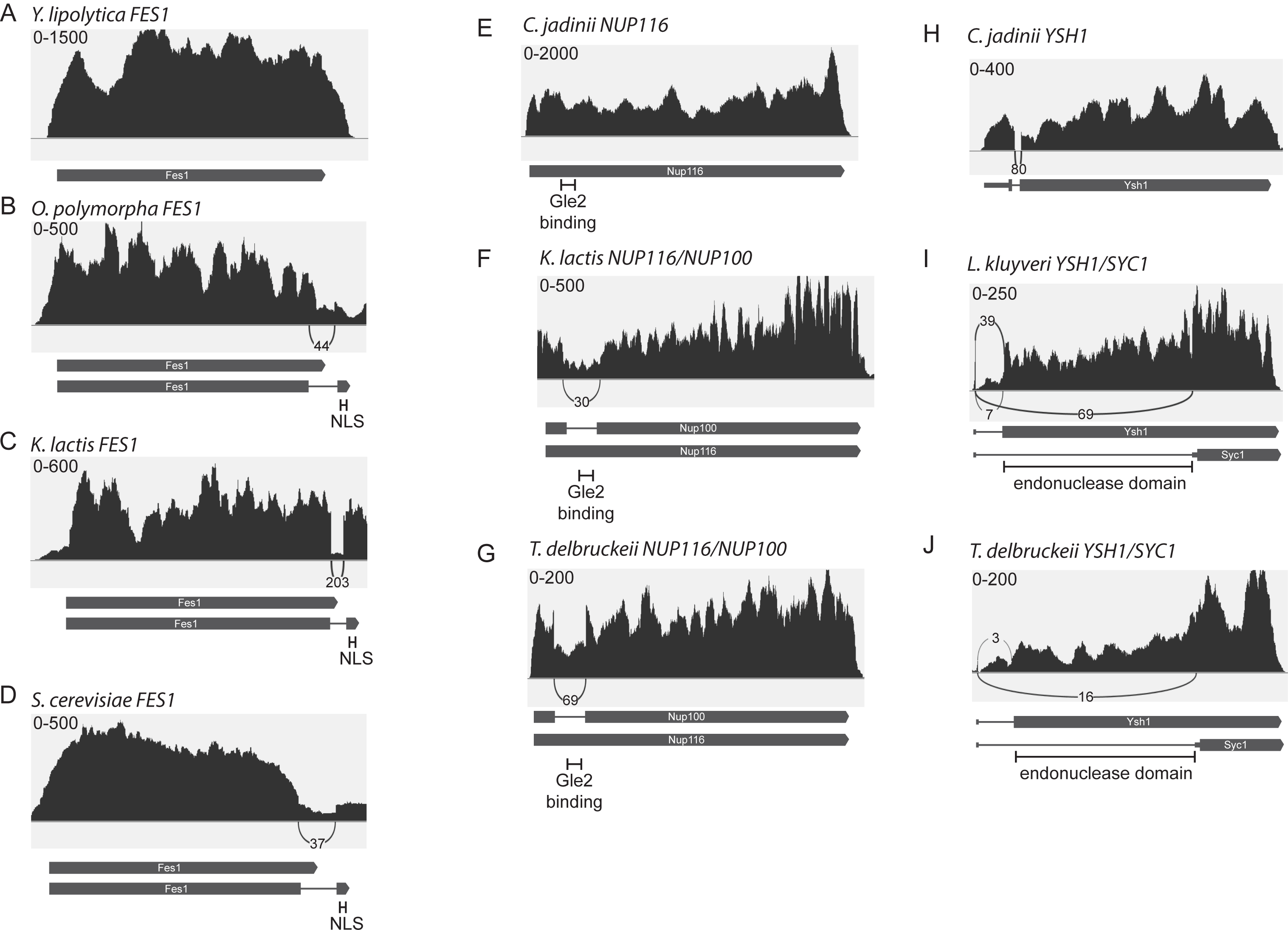
Conservation of alternative splicing patterns during divergence of Saccharomycotina. All panels show Sashimi plots from representative RNA-seq data sets. Numbers in the top left corner indicate the scale of the y-axis in number of reads. Numbers next to arches indicate the number of exon junction reads across the intron. The encoded proteins are depicted below the plots. **A**: The *Y. lipolytica FES1* gene is not spliced. **B, C, and D:** The *FES1* genes of *O. polymorpha, K. lactis* and *S. cerevisiae* use intron retention to encode a cytoplasmic Fes1 and splicing to include a conserved NLS that in *S. cerevisiae* has been shown to target the protein to the nucleus. **E**: The *C. jadinii NUP116* gene is not spliced. **F and G:** Saccharomycetaceae of the KLE (panel F) and ZT (panel G) clade use intron retention to encode Nup116 and Nup100 from a single gene. Indicated is the conserved sequence that has previously been show to act as a binding site for Gle2 in Nup116 and distinguishes Nup116 function from Nup100. **H:** The *C. jadinii YSH1* gene contains a constitutively spliced intron between codons 4 and 5. **I and J:** Saccharomycetaceae of the KLE (panel I) and ZT (Panel J) clades use alternative 3’ splice sites to encode Ysh1-like and Syc1-like proteins from a single gene. Indicated is the conserved endonuclease domain of Ysh1 that is absent from Syc1. *L. kluyveri* uses two different proximal 3’ splices site upstream of the start codon for Ysh1 and a distal 3’ splice site upstream of the start codon for Syc1.

Overall, these results indicate that all of the alternative splicing events studied here arose over 100 MYA and are each conserved in multiple families. These conservation times are similar to the divergence of vertebrates from each other. For example, human and fish diverged an estimated 435 MYA, while humans diverged from marsupials 159 MYA and from mice 90 MYA. Our observation that these alternative splicing events are conserved for more than 100 million years further confirms that they are functionally important.

### Alternative splicing arose by intronization and exonization

Alternative splicing can arise by generation of an intron from previously exonic events (intronization), or by generating an exon out of previously intronic sequences (exonization). *NUP116/NUP100* and *YSH1/SYC1* provide clear examples of intronization: exonic sequences that were converted to introns. The functional difference between Nup116 and Nup100 is a Gle2 binding site that is present in Nup116 but absent in Nup100 (Bailer et al., 1998). This sequence is conserved in the retained intron of the KLE and ZT clades (Figure 2F and G) but more importantly, even in the orthologs that diverged before alternative splicing arose, such as *C. jadinii* (Figure 2E). This indicates that the Gle2-interacting sequence existed before alternative splicing arose and was lost in Nup100 by converting the encoding exon into an intron. The intron was then lost post-WGD to form the current *Saccharomyces NUP100* gene. Similarly, Ysh1 has an N-terminal endoribonuclease domain that functions in 3’ end formation of mRNAs through cleavage and polyadenylation, and a C-terminal protein-protein interaction domain (Lidschreiber et al., 2018; Nedea et al., 2003). This N-terminal domain is clearly conserved in the orthologs that diverged before alternative splicing arose (Figure 2H; and even in Metazoa). Some orthologs that diverged before alternative splicing arose contain a constitutive intron close to the 5’ end of the mRNA (Figure 2H). Alternative splicing appears to have arisen by the evolution of an additional 3’ splice site within the coding region that causes the entire endonuclease domain to be skipped in the Syc1-like protein (Figure 2 I and J. This addition of a distal 3’ splice site converted a (constitutive) exon to an (alternative) intron.

We detected one example of exonization: the conversion of intronic sequences to exonic sequences in the *SRC1* gene of the ZT clade. The *SRC1* gene contains anciently conserved 5’ splice sites to encode two different proteins (Figure 3A). In addition, in the ZT clade all the stop codons were lost from the intron (Figure 3C). Therefore, a single *SRC1/HEH2* gene in the ZT clade encodes three proteins: one through intron retention and the other two through alternative 5’ splice sites (Figure 3C).

**Figure 3:**
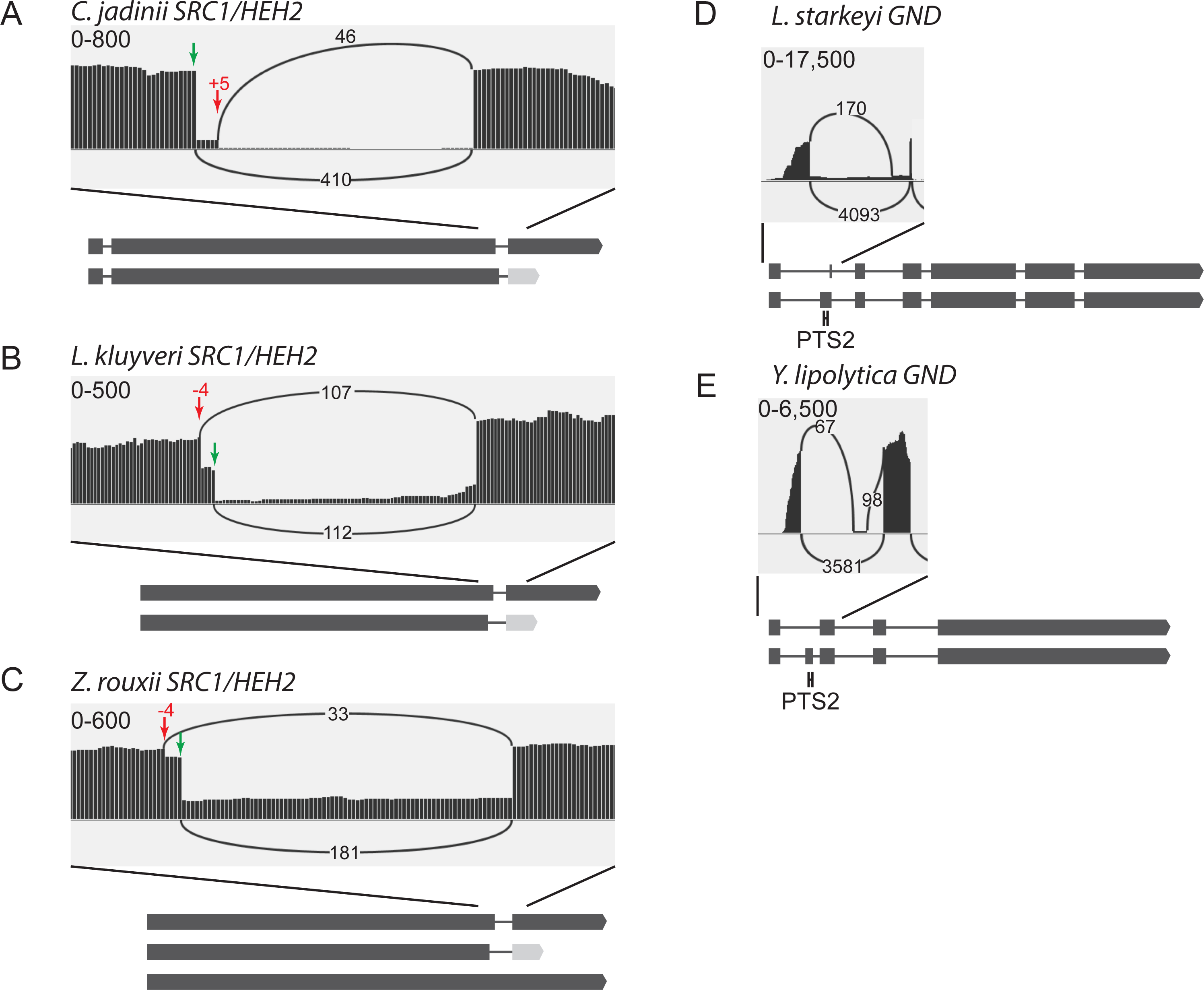
Changes in alternative splicing patterns during divergence of Saccharomycotina. All panels show Sashimi plots from representative RNA-seq data sets. Numbers in the top left corner indicate the scale of the y-axis in number of reads. Numbers next to arches indicate the number of exon junction reads across the intron. The encoded proteins are depicted below the plots. **A**: Early diverging Saccharomycotina, including *C. jadinii* encode a truncated *Src1/Heh2* protein by using an alternative 5’ splice site (red arrow) that is 5 nucleotides distal to the splice site used for the full-length protein (green arrow). **B:** The alternative 5’ splice site is shifted to −4 in the Saccharomycetaceae. **C:** in the ZT clade the ortholog encodes an additional intron retention isoform. **D:** Top panel: *L. starkeyi* uses alternative 3’ splice sites in intron 1 to either include or skip a PTS2 sorting signal. **E:** *Y. lipolytica* uses inclusion or skipping of exon 2 to either include or skip a PTS2 sorting signal.

In contrast to these clear examples of intronization in *YSH1/SYC1* and *NUP116/NUP100* and exonization in *SRC1/HEH2*, we were not able to clearly establish the origin of the retained intron sequences of *PTC7* and *FES1*.

### Loss of alternative splicing events often follows gene duplication

Our RNA-seq analysis revealed ten instances where an alternative splicing event was lost in the Saccharomycotina. The parents of the WGD event appear to have used alternative splicing in six different genes. Five of these six genes are replaced by a pair of duplicated genes in at least some WGD species, with three different fates. In two gene pairs (*SKI7/HBS1* and *NUP116/NUP100*) both paralogs lost alternative splicing in all four WGD species examined, such that the two isoforms are now expressed from different paralogs. In two genes (*PTC7* and *YSH1/SYC1*) some WGD species maintained alternative splicing of one paralog and lost the other paralog, while in other WGD species the two isoforms are now expressed from two different paralogs. Specifically one *T. blattae PTC7* ortholog (*TBLA_0A03190*) is constitutively spliced and predicted to encode a mitochondrial protein, while the other ortholog (*TBLA_0H01920*) lost the capacity to be spliced and is predicted to encode a protein inserted into the nuclear envelope. These different outcomes in different WGD species suggests that the loss of alternative splicing does not always take place soon after duplication, but can take place after sufficient time has elapsed for speciation.

The loss of alternative splicing of the *SRC1/HEH2* gene pair is complex. As detailed above, this gene encodes two full-length proteins and a truncated protein in the ZT clade, and one full-length and one truncated protein in the KLE clade. The ancestral WGD species thus likely inherited the capacity to make five distinct proteins, but each extant WGD species only maintained three, with one gene (*SRC1*) encoding two proteins by the use of 5’ splice sites (or in *Naumovozyma castellii* through the retained intron) and the other gene (*HEH2*) encoding the third protein. Thus, while casual inspection suggests that all four WGD species have maintained alternative splicing of *SRC1* and added an *HEH2* gene, closer inspection suggest that the immediate product of WGD encoded five Src1/Heh2 proteins and each WGD species has maintained three of these proteins. Since the three proteins are never expressed from the same gene, it appears that in this case duplicate genes also replaced alternative splicing. A better understanding of the functional differences between Heh2 and Src1 is required to completely understand their evolutionary history. One possibility arising from our analysis is that the evolution of intron retention in the ZT clade necessitated the retention of three different proteins in the WGD clade.

The only alternatively spliced gene in the WGD clade that was not replaced by a pair of genes is *FES1*. Three species have maintained the alternative splicing event, while *T. blattae* has lost the intron, but has not retained a duplicate gene. Thus, *T. blattae* appears to have retained only the cytoplasmic unspliced isoform. Interestingly, the same loss occurred in the lineage leading to *C. albicans* and *C auris.* Therefore, either *T. blattae, C. albicans* and *C. auris* have lost nuclear Fes1 or they use an alternative mechanism to target this Hsp70 nucleotide exchange factor to the nucleus. Overall, our analyses indicate that WGD was frequently followed by loss of alternative splicing.

Interestingly, loss of alternative splicing is more frequent in WGD species than other Saccharomycotina. Specifically, the WGD clade diverged at about the same time that *C. albicans* diverged from *C. auris, K. lactis* from *L. kluyveri* and *Z. rouxii* from *T. delbrueckii* (Figure 1), but only one loss of an alternative splicing event occurred in any of these six species (*YSH1/SYC1* in *Z. rouxii*; Figure 1).

While loss of alternative splicing is more common after WGD, we detected four losses that did not follow the WGD, of which two were accompanied by small-scale duplications. First, it has previously been shown that the *Y. lipolytica* malate dehydrogenase (MDH) gene uses alternative 3’ splice sites (Kabran et al., 2012). When the distal 3’ splice site is used, the protein includes a PTS1 (Peroxisomal Targeting Signal 1) and is targeted to the peroxisomes (Kabran et al., 2012), similar to Mdh3 in *Saccharomyces*. When the proximal 3’splice site is used, the encoded protein remains in the cytoplasm (Kabran et al., 2012), similar to Mdh2 in *Saccharomyces*. This single alternatively spliced MDH gene was replaced with duplicate *MDH2* and *MDH3* genes between 304 and 322 MYA in a common ancestor of *Candida* and *Saccharomyces*. One of the duplicate genes in each species encodes a PTS1 signal and the encoded protein is predicted peroxisomal. Specifically, alternative splicing results in a C-terminal AKI or AKM (in *Y. lipolytica* and *L. starkeyi*, respectively), while one of the duplicated genes in the other species encodes a C-terminal S/T/A, K, L/I/M. The other gene lacks any resemblance to a PTS1 signal and is predicted to be cytoplasmic. Second, in *O. polymorpha* the alternatively spliced *SRC1/HEH2* gene is replaced by duplicated genes that now each express one of the isoforms (the full-length and truncated forms mentioned above). Third, it has previously been shown that the *C. albicans* 6-phosphogluconate dehydrogenase (GND) gene uses alternative 3’ splice sites (Strijbis et al., 2012). In this case, use of the proximal 3’ splice site causes inclusion of a PTS2 signal and targeting to the peroxisome, while use of the distal 3’ splice site results in a cytoplasmic protein (Strijbis et al., 2012). We observed this alternative splicing in early diverging species (Figure 1) but this appears to have been lost twice, in *O. polymorpha* and in the Saccharomycetaceae (both >114 MYA). As a result both *O. polymorpha* and the Saccharomycetaceae may have lost the peroxisomal isoform. Overall this suggests that most losses of alternative splicing in the Saccharomycotina were accompanied by either WGD or small-scale duplications, thereby preserving the functional divergence of the proteins.

### Changes in alternative splicing events

In addition to gains and losses of alternative splicing, we also detected six changes in alternative splicing pattern. First, we have previously used rt-PCR and RACE to show that in *C. albicans SKI7/HBS1* the use of the proximal 3’ splice site is replaced by the use of an alternative promoter that is located inside the intron (Marshall et al., 2013). This was confirmed by RNA-seq data. Second, the ancestral Saccharomycotina used an alternative splice site in the *SRC1/HEH2* that was 5 nucleotides distal to the canonical site (Figure 3A). This alternative site was shifted nine nucleotides from +5 to −4 of the canonical splice site in an early Saccharomycetaceae (Figure 3B). Third, this same change from +5 to −4 occurred in *C. albicans SCR1/HEH2*. Because *C. auris* and *C. jadinii* maintain the ancestral alternative 5’ splice site at +5, the most parsimonious explanation is that independent changes occurred in Saccharomycetaceae and *C. albicans*. Fourth, *A. fumigatus* uses alternative 3’ splices sites in the orthologous intron of *SRC1/HEH2* instead of alternative 5’ splice sites, suggesting that another shift occurred either very early in the Saccharomycotina or in the *Aspergillus* lineage. Fifth, several species use alternative 3’ splice site to include a PTS2 in *GND1/GND2* (Figure 3D; (Strijbis et al., 2012)) while *Y. lipolytica* has switched to exon skipping (Figure 3E). Sixth, *A. fumigatus* uses both intron retention (leading to an alternative stop codon) and alternative 3’ splice sites for its *MDH2/MDH3* gene. *L. starkeyi* uses only the retention mechanism, and *Y. lipolytica* only the alternative 3’ splice sites. While the splicing signals and patterns change in each of these cases, protein alignments suggest that the functional consequences are conserved.

## Conclusions

Here, we survey the evolution of alternative splicing in the Saccharomycotina during their 330 million year divergence. Although alternative splicing is rarer in the Saccharomycotina than in Metazoa, all 14 Saccharomycotina species studied here use alternative splicing to functionally diversify their proteome and all eight events that we have characterized here are conserved for at least 100 million years, and several for 500 million years. On the other hand, no single gene is alternatively spliced in all 14 species. There are likely additional alternative splicing events in these species, but we have limited our analysis to events that had clear and previously described functional consequences (See Table 1). Loss of alternative splicing seems to often be mediated by a gene duplication that maintains the functional diversification. We anticipate that other small-scale duplication and WGD events, including the two rounds of WGD in the human lineage had similar effects.

## Materials and methods

The times of species divergence cited in the text and in Figure 1 are from timetree.org and represent a consensus of available data.

We performed extensive literature searches for “alternative splicing” and each of our target species (Table 1). We included all examples with some evidence that the alternative splicing event resulted in functionally distinct proteins. The previously predicted alternative splicing of PGK in *Y. lipolytica* (Freitag et al., 2012) could not be detected in RNA-seq data from that species (data not shown) and thus was not included. We also excluded some cases of regulated splicing where only the amount of the encoded protein is affected because this would require quantitative RNA-seq analysis from many different conditions (e.g. *C. albicans DUR31* (Donovan et al., 2018) and *S. cerevisiae MTR2* (Davis et al., 2000), *GCR1* (Hossain et al., 2016) and *APE2* (Meyer et al., 2011)). This literature search was supplemented by inspecting RNA-seq data from two strategically placed species, *L. kluyveri* and *C. albicans*, which identified one novel intron retention event (*L. kluyveri SAKL0B11770g*, corresponding to the WGD paralogs *NUP100* and *NUP116* in *S. cerevisiae*).

For most species, raw RNA-seq reads from previous studies RNA-seq data were downloaded from EBI (https://www.ebi.ac.uk/ena). Because there was no publicly available RNA-seq data from the *Tetrapisispora/Vanderwaltozyma* branch we generated RNA-seq data from *T. blattae* (deposited in SRA under project PRJNA613484). As explained above, this branch diverged soon after WGD and thus was critical in understanding the fate of alternative splicing after WGD. We obtained *T. blattae* from the United States Department of Agriculture, Agricultural Research Service Culture Collection (https://nrrl.ncaur.usda.gov/), grew duplicate cultures in YPD medium at 30 °C, isolated RNA by a hot phenol method (He et al., 2008) and generated paired-end 150 nt RNA-seq data (25,338,942 and 30,978,862 pairs for the biological replicates).

Both downloaded and newly generated RNA-seq reads were adaptor and quality trimmed with Trim Galore when needed (https://www.bioinformatics.babraham.ac.uk/projects/trim_galore/), and aligned to the genome downloaded from Genbank (https://www.ncbi.nlm.nih.gov/genome/) using TopHat2 (Kim et al., 2013). Several of the datasets were also aligned with HiSat2 (Kim et al., 2015) and RNA STAR (Dobin et al., 2013) with identical results. At least two datasets from each species were analyzed (Supplemental Table 1). If more datasets were available, preference was given to datasets generated in two independent studies of the same species, paired-end reads, 150 nt reads, datasets with 20-30 million reads, datasets from the same wild-type strain as that used for whole genome sequencing and datasets from standard growth conditions. These preferences were all designed to ease the detection of alternative splicing. TopHat2 settings were adjusted to allow introns from 30 nucleotides to 10 kb in size. The resulting alignment files were manually inspected in IGV (https://software.broadinstitute.org/software/igv/), and Sashimi plots were generated in IGV.

A gene was considered to use alternative splice sites if the number of exon junction reads for the minor splicing patterns was at least 10% of that for the major splicing pattern detected. An exception was made for *GND1/GND2* alternative splicing because it has previously been shown that the PTS2-included form accounts for only ∼0.1% of the mRNA, yet is functionally important (Strijbis et al., 2012). This low level of alternative splicing could be reliably detected because of the very high expression of this gene (Figure 3D and E). The *PTC7* and *NUP100/NUP116* intron was considered retained if the read coverage within the intron was at least 10% of the exon junction reads, if the intron length was a multiple of 3, and if the intron contained no in frame stop codons. For *MDH2/MDH3* and *FES1* the intron was considered to be retained if read coverage within the intron was at least 10% of the exon junction reads, and extended past the first in-frame stop codon.

To understand the functional divergence of splice isoforms and duplicate genes, multiple sequence alignments were generated with Clustal Omega (https://www.ebi.ac.uk/Tools/msa/clustalo/) after correcting some of the protein sequences based on the observed splicing patterns. PSORT II (https://psort.hgc.jp/form2.html) was used to aid in the prediction of protein localization, and TMHMM2.0 was used to predict transmembrane helices in *PTC7* http://www.cbs.dtu.dk/services/TMHMM/.

## Acknowledgements

This work was supported by NIH grant R01GM099790 to AvH

## References

Araki, Y., Takahashi, S., Kobayashi, T., Kajiho, H., Hoshino, S., and Katada, T. (2001). Ski7p G protein interacts with the exosome and the Ski complex for 3’-to-5’ mRNA decay in yeast. EMBO J 20, 4684–4693.

Bailer, S.M., Siniossoglou, S., Podtelejnikov, A., Hellwig, A., Mann, M., and Hurt, E. (1998). Nup116p and nup100p are interchangeable through a conserved motif which constitutes a docking site for the mRNA transport factor gle2p. EMBO J 17, 1107–1119.

Becker, T., Armache, J.P., Jarasch, A., Anger, A.M., Villa, E., Sieber, H., Motaal, B.A., Mielke, T., Berninghausen, O., and Beckmann, R. (2011). Structure of the no-go mRNA decay complex Dom34-Hbs1 bound to a stalled 80S ribosome. Nature structural & molecular biology 18, 715–720.

Davis, C.A., Grate, L., Spingola, M., and Ares, M., Jr. (2000). Test of intron predictions reveals novel splice sites, alternatively spliced mRNAs and new introns in meiotically regulated genes of yeast. Nucleic Acids Res 28, 1700–1706.

Dobin, A., Davis, C.A., Schlesinger, F., Drenkow, J., Zaleski, C., Jha, S., Batut, P., Chaisson, M., and Gingeras, T.R. (2013). STAR: ultrafast universal RNA-seq aligner. Bioinformatics 29, 15–21.

Donovan, P.D., Holland, L.M., Lombardi, L., Coughlan, A.Y., Higgins, D.G., Wolfe, K.H., and Butler, G. (2018). TPP riboswitch-dependent regulation of an ancient thiamin transporter in Candida. PLoS Genet 14, e1007429.

Freitag, J., Ast, J., and Bolker, M. (2012). Cryptic peroxisomal targeting via alternative splicing and stop codon read-through in fungi. Nature 485, 522–525.

Gowda, N.K., Kaimal, J.M., Masser, A.E., Kang, W., Friedlander, M.R., and Andreasson, C. (2016). Cytosolic splice isoform of Hsp70 nucleotide exchange factor Fes1 is required for the degradation of misfolded proteins in yeast. Mol Biol Cell 27, 1210–1219.

Grund, S.E., Fischer, T., Cabal, G.G., Antunez, O., Perez-Ortin, J.E., and Hurt, E. (2008). The inner nuclear membrane protein Src1 associates with subtelomeric genes and alters their regulated gene expression. The Journal of cell biology 182, 897–910.

He, F., Amrani, N., Johansson, M.J., and Jacobson, A. (2008). Chapter 6. Qualitative and quantitative assessment of the activity of the yeast nonsense-mediated mRNA decay pathway. Methods Enzymol 449, 127–147.

Hossain, M.A., Claggett, J.M., Edwards, S.R., Shi, A., Pennebaker, S.L., Cheng, M.Y., Hasty, J., and Johnson, T.L. (2016). Posttranscriptional Regulation of Gcr1 Expression and Activity Is Crucial for Metabolic Adjustment in Response to Glucose Availability. Mol Cell 62, 346–358.

Juneau, K., Nislow, C., and Davis, R.W. (2009). Alternative splicing of PTC7 in Saccharomyces cerevisiae determines protein localization. Genetics 183, 185–194.

Kabran, P., Rossignol, T., Gaillardin, C., Nicaud, J.M., and Neuveglise, C. (2012). Alternative splicing regulates targeting of malate dehydrogenase in Yarrowia lipolytica. DNA research : an international journal for rapid publication of reports on genes and genomes 19, 231–244.

Kellis, M., Birren, B.W., and Lander, E.S. (2004). Proof and evolutionary analysis of ancient genome duplication in the yeast Saccharomyces cerevisiae. Nature 428, 617–624.

Kim, D., Langmead, B., and Salzberg, S.L. (2015). HISAT: a fast spliced aligner with low memory requirements. Nat Methods 12, 357–360.

Kim, D., Pertea, G., Trapnell, C., Pimentel, H., Kelley, R., and Salzberg, S.L. (2013). TopHat2: accurate alignment of transcriptomes in the presence of insertions, deletions and gene fusions. Genome Biol 14, R36.

Kowalinski, E., Kogel, A., Ebert, J., Reichelt, P., Stegmann, E., Habermann, B., and Conti, E. (2016). Structure of a Cytoplasmic 11-Subunit RNA Exosome Complex. Mol Cell 63, 125–134.

Lidschreiber, M., Easter, A.D., Battaglia, S., Rodriguez-Molina, J.B., Casanal, A., Carminati, M., Baejen, C., Grzechnik, P., Maier, K.C., Cramer, P., et al. (2018). The APT complex is involved in non-coding RNA transcription and is distinct from CPF. Nucleic Acids Res 46, 11528–11538.

Marcet-Houben, M., and Gabaldon, T. (2015). Beyond the Whole-Genome Duplication: Phylogenetic Evidence for an Ancient Interspecies Hybridization in the Baker’s Yeast Lineage. PLoS Biol 13, e1002220.

Marshall, A.N., Han, J., Kim, M., and van Hoof, A. (2018). Conservation of mRNA quality control factor Ski7 and its diversification through changes in alternative splicing and gene duplication. Proc Natl Acad Sci U S A 115, E6808–E6816.

Marshall, A.N., Montealegre, M.C., Jimenez-Lopez, C., Lorenz, M.C., and van Hoof, A. (2013). Alternative splicing and subfunctionalization generates functional diversity in fungal proteomes. PLoS genetics 9, e1003376.

Meyer, M., Plass, M., Perez-Valle, J., Eyras, E., and Vilardell, J. (2011). Deciphering 3’ss selection in the yeast genome reveals an RNA thermosensor that mediates alternative splicing. Mol Cell 43, 1033–1039.

Nedea, E., He, X., Kim, M., Pootoolal, J., Zhong, G., Canadien, V., Hughes, T., Buratowski, S., Moore, C.L., and Greenblatt, J. (2003). Organization and function of APT, a subcomplex of the yeast cleavage and polyadenylation factor involved in the formation of mRNA and small nucleolar RNA 3’-ends. The Journal of biological chemistry 278, 33000–33010.

Pisareva, V.P., Skabkin, M.A., Hellen, C.U., Pestova, T.V., and Pisarev, A.V. (2011). Dissociation by Pelota, Hbs1 and ABCE1 of mammalian vacant 80S ribosomes and stalled elongation complexes. The EMBO journal 30, 1804–1817.

Rodriguez-Navarro, S., Igual, J.C., and Perez-Ortin, J.E. (2002). SRC1: an intron-containing yeast gene involved in sister chromatid segregation. Yeast 19, 43–54.

Scannell, D.R., Frank, A.C., Conant, G.C., Byrne, K.P., Woolfit, M., and Wolfe, K.H. (2007). Independent sorting-out of thousands of duplicated gene pairs in two yeast species descended from a whole-genome duplication. Proceedings of the National Academy of Sciences of the United States of America 104, 8397–8402.

Shen, X.X., Zhou, X., Kominek, J., Kurtzman, C.P., Hittinger, C.T., and Rokas, A. (2016). Reconstructing the Backbone of the Saccharomycotina Yeast Phylogeny Using Genome-Scale Data. G3 (Bethesda) 6, 3927–3939.

Shoemaker, C.J., Eyler, D.E., and Green, R. (2010). Dom34:Hbs1 promotes subunit dissociation and peptidyl-tRNA drop-off to initiate no-go decay. Science 330, 369–372.

Strijbis, K., van den Burg, J., Visser, W.F., van den Berg, M., and Distel, B. (2012). Alternative splicing directs dual localization of Candida albicans 6-phosphogluconate dehydrogenase to cytosol and peroxisomes. FEMS yeast research 12, 61–68.

van Hoof, A., Staples, R.R., Baker, R.E., and Parker, R. (2000). Function of the ski4p (Csl4p) and Ski7p proteins in 3’-to-5’ degradation of mRNA. Mol Cell Biol 20, 8230–8243.

